# Analysis of Long-Term Neuronal Dynamics via Ordinal Pattern Quantifiers Following Traumatic Brain Injury and Pharmacological Modulation

**DOI:** 10.64898/2026.04.30.721848

**Authors:** Maria Moro-Fernández, Alejandro Carretero-Guillén, Jon Ondaro, Xabier Bengoetxea, Inés Moreno-Jiménez, Roger Pradés, Juan Manuel Encinas-Pérez, Diego M. Mateos

**Author notes:** These authors contributed equally (Maria Moro-Fernández, Alejandro Carretero-Guillén).

## Abstract

Traumatic brain injury (TBI) profoundly disrupts hippocampal network dynamics, triggering persistent alterations in oscillatory activity that underlie cognitive deficits and increased susceptibility to post-traumatic epilepsy. Characterizing these alterations quantitatively remains challenging: the resulting signals are nonlinear, non-stationary, and exhibit complex multiscale structure that conventional spectral metrics fail to resolve. Ordinal-pattern information-theoretic quantifiers offer a principled, model-free alternative for probing such dynamics. In this work we apply permutation entropy (PE), statistical complexity (SC), Fisher information (FI), and permutation Lempel–Ziv complexity (PLZC) to hippocampal local field potentials (LFPs) recorded over 21 days in a rodent controlled cortical impact model of TBI, across five experimental groups under distinct pharmacological conditions. Embedding signal trajectories in the SC–PE, FI–PE, and PLZC–PE information planes reveals group- and time-dependent dynamical signatures in the theta (4–8 Hz) and high-frequency oscillation (80–200 Hz) bands, exposing state transitions invisible to spectral analysis. Unsupervised dimensionality reduction (UMAP) combined with HDBSCAN clustering further delineates distinct regions of dynamical state space associated with injury progression and pharmacological modulation. We additionally applied the Ordinal Modulation Index (OMI), an ordinal-based measure of theta–HFO cross-frequency coupling, which captures treatment-dependent reorganization of phase–amplitude coordination. These results establish ordinal-pattern analysis as a sensitive and interpretable framework for tracking the nonlinear reorganization of hippocampal dynamics following TBI.

## 1 Introduction

Traumatic brain injury (TBI) is an alteration in brain function caused by an external mechanical force—such as a blow, blast, or penetrating impact to the head—that disrupts normal neuronal activity through a combination of primary mechanical damage and a cascade of secondary pathophysiological processes, including neuroinflammation, excitotoxicity, axonal injury, and progressive cell death [1, 2]. From an epidemiological standpoint, TBI represents a major global health burden: the Global Burden of Disease Study reported approximately 27 million new cases and nearly 49 million prevalent cases worldwide in 2019, contributing to over 7 million years lived with disability [2]. Beyond acute mortality, survivors frequently endure persistent cognitive, behavioral, and neurological sequelae that impose substantial personal and societal costs [2].

The hippocampus is especially susceptible to TBI at the neural circuit level. Experimental models consistently demonstrate selective neuronal loss in the CA3 subfield and the dentate hilus, alongside progressive mossy-fiber sprouting and bilateral hyperexcitability in the dentate gyrus, reflecting the structural characteristics of hippocampal sclerosis [3]. These pathological alterations disturb the dynamics of hippocampal local field potential (LFP): research indicates a reduction in theta (4–8 Hz) oscillatory power, compromised theta–gamma phase–amplitude coupling (PAC), diminished sharp-wave ripple amplitudes, and decreased entrainment of pyramidal cells and interneurons to synchronized oscillatory activity [4, 5, 6].

A clinically significant outcome of TBI-induced hippocampal hyperexcitability is the development of post-traumatic epilepsy (PTE). Epidemiological studies indicate that traumatic brain injury (TBI) is the discernible cause of epilepsy in approximately 30% individuals aged 15–34 who experience seizures [7], with cumulative incidence rates reaching 25% at five years and 32% fifteen years post-severe TBI [8]. At the same time, pathological high-frequency oscillations (HFOs, 80–500 Hz) develop in the hippocampus post-TBI; these phenomena imply hypersynchronous network states and have been recognized as biomarkers of epileptogenesis and increased seizure susceptibility [9, 10].

The current standard of care for post-traumatic seizure prophylaxis primarily involves anti-seizure medications like levetiracetam (LEV), which is extensively utilized to mitigate early seizures due to its advantageous pharmacokinetics and comparable efficacy to phenytoin. However, LEV functions merely as a symptomatic treatment, failing to interrupt the underlying epileptogenic process, thereby leaving late seizures and long-term cognitive deficits unaddressed [11, 12].

In this study, we examine how hippocampal baseline dynamics evolve over time in rats subjected to controlled cortical impact (CCI), a well-established rodent model of TBI, with and without pharmacological treatment, relative to uninjured sham controls. Levetiracetam (LEV) was administered as the treatment regimen for 21 days following injury. We recorded LFPs over four weeks, five months before treatment, and looked at the theta (4–8 Hz) and HFO (80–200 Hz) frequency bands. We chose these bands because theta oscillations help the hippocampus do memory-related calculations [5] and HFOs are sensitive markers of network hyperexcitability and epileptogenesis [9].

Furthermore, we differentiate from traditional spectral methodologies and instead measure hippocampal LFP dynamics using an information-theoretic framework grounded in ordinal-pattern symbolization [13]. We specifically calculate Permutation Entropy (PE) [13], Statistical Complexity (SC)[14], Fisher information (FI) [15], and Permutation Lempel–Ziv Complexity (PLZC) [16] from the ordinal-pattern probability distribution of each signal. This group of quantifiers works especially well with neural time series because they are fast to compute, resistant to noise, don’t change when the data is transformed in a monotonic way, and are sensitive to changes in the structure of time without needing to make assumptions about stationarity [17]. The complexity—entropy and Fisher—entropy planes generated by these measures offer a concise and comprehensible assessment of a signal’s proximity to stochastic noise, deterministic chaos, or structured periodic dynamics. To further explore the high-dimensional structure of these ordinal-pattern feature spaces, we apply Uniform Manifold Approximation and Projection (UMAP) [18] combined with HDBSCAN clustering [19], enabling an unsupervised assessment of whether experimental groups occupy distinct regions in the information-theoretic feature space. As far as we know, this is the first time that ordinal-pattern information quantifiers have been used to look at LFP dynamics in a TBI model.

We also use the Ordinal Modulation Index (OMI) [20], a new version of the classical modulation index [21], to measure cross-frequency coupling between theta phase and HFO amplitude. This adds to single-band analyses. This method involves discretizing the HFO amplitude time series using ordinal patterns before calculating the coupling measure. The OMI offers a logical extension of the ordinal-pattern framework to cross-frequency interactions by combining the noise robustness of ordinal symbolization with the sensitivity of phase–amplitude coupling analysis.

These analyses characterize how TBI alters the information-theoretic structure of hippocampal LFP dynamics across frequency bands, and whether LEV administration restores or modifies this structure in the weeks following injury.

## 2 Methods

### 2.1 Animal Model and Experimental Groups

Experiments were performed on adult male and female C57BL/6J mice (12 weeks of age; Janvier Labs, France). Traumatic brain injury was induced by controlled cortical impact (CCI) following a previously described protocol [22]. All experimental procedures were approved by the University of the Basque Country (EHU/UPV) Ethics Committee (Leioa, Spain) and the Comunidad Foral de Bizkaia (CEEA: M20/2015/236).

Sham-operated animals underwent identical surgical procedures without impact. Following injury, animals were assigned to three experimental groups: SHAM (*n* = 5), TBI-SAL (*n* = 20), and TBI-LEV (*n* = 16). Levetiracetam was administered subcutaneously once daily for 21 days beginning 45 minutes post-injury. Five months after TBI, animals were implanted with intracerebral electrodes using stereotaxic coordinates from a standard rat brain atlas [23].

### 2.2 LFP Recordings

In vivo LFP recordings were obtained five months after TBI or sham surgery. Animals were anesthetized with isoflurane (3–4% induction, 1–2% maintenance) and placed in a stereotaxic frame for electrode implantation. A small craniotomy was made above the dorsal hippocampus, and a tungsten microelectrode (50 µm diameter) was lowered into the dentate gyrus using standard stereotaxic coordinates. A cerebellar silver wire served as a reference and ground. Electrodes were fixed with dental acrylic, and animals recovered for at least one week before recordings. LFPs were acquired in awake, freely moving mice inside an open-field recording chamber, amplified and bandpass filtered (1–500 Hz), and digitized at 10 kHz using an Intan-based system. Electrode placement was confirmed histologically. This preparation enabled stable long-term monitoring of hippocampal dynamics for ordinal pattern analysis.

### 2.3 Signal Preprocessing

Raw LFP signals were downsampled from 10.000 Hz to 1.000 Hz to reduce computational load while preserving the frequency content of interest. The signals were band-pass filtered between 1 and 500 Hz and notch-filtered at 50, 100, 150, and 200 Hz to remove line noise and its harmonics.

The preprocessed signals were segmented into non-overlapping 10 s windows and subjected to an automatic quality-control procedure. For each segment, power spectral density was estimated using Welch’s method, and a frequency-domain signal-to-noise ratio (SNR_freq_) was computed as the ratio between mean power in the 1–200 Hz band and mean power above 200 Hz. Segments with SNR_freq_ < 15 were rejected. A robust time-domain SNR (SNR_time_) was calculated as the ratio between the median absolute amplitude and the interquartile range; segments with SNR_time_ < 0.5 were discarded. Segments with variance exceeding the mean variance across segments from the same channel were also excluded. In addition, the Hurst exponent was estimated via log–log regression of lagged differences, and only segments with Hurst values within the 10th–90th percentile range of the channel-specific distribution were retained. Segments satisfying all criteria were considered valid for subsequent analyses (see Supplementary Section S1 for details).

After quality control, recordings with fewer than 30 valid epochs were not used in subsequent analyses. The average number of retained epochs across the recordings that were looked at was N_*epoch*_ = 285.4 ± 203.7 (range: [52, 880]).

### 2.4 Analysis

#### 2.4.1 Frequensy bands

To identify potential treatment-related variations in brain dynamics, we focused on two frequency bands—theta (4–8 Hz) and high-frequency oscillations (HFOs, 80–250 Hz)—along with their cross-frequency coupling.

HFOs, typically classified as ripples, have emerged as reliable biomarkers of epileptogenic tissue and are consistently observed in both ictal and interictal recordings across epilepsy patients and animal models [24]. Critically, their spatial distribution and rate correlate with seizure onset zones and postoperative outcomes [25], making them a particularly informative readout of residual epileptogenic activity at chronic time points. In the context of post-traumatic epilepsy, the persistence or suppression of aberrant high-frequency oscillatory activity at six months may indicate early LEVE treatment may have effectively interrupted the underlying epileptogenic cascade or postponed its advancement.

The theta band, in turn, provides a complementary perspective at the network level. Theta oscillations are closely associated with hippocampal–cortical communication and large-scale network synchronization, and their dynamics are known to be disrupted in epileptic networks and during epileptogenesis [26, 27, 28]. Theta-band activity at the chronic stage provides a window into whether early pharmacological intervention has preserved, changed, or restored network-level function because TBI can cause long-term structural and functional reorganization of hippocampal circuits.

Beyond their individual relevance, theta and HFO activity are functionally linked through cross-frequency interactions. The hierarchical structure of neural circuits is reflected in phase-amplitude coupling (PAC) between low-frequency oscillations and high-frequency activity, where slow rhythms coordinate local high-frequency processing across larger networks [29]. This coupling may become abnormal under pathological circumstances, indicating network hypersynchrony and a disturbed excitation-inhibition balance [25, 30].

#### 2.4.2 Ordinal Patterns

Ordinal patterns quantify the temporal structure of a time series by analyzing the relative ordering of neighboring values rather than their absolute amplitudes [13].

Given a time series {*x*_*t*_} of length *N*, an embedding dimension *d* ≥ 2, and a time delay *τ*, each point *x*_*t*_ is mapped to an ordinal pattern *π* ∈ Π by ranking the *d* successive values (*x*_*t*_, *x*_*t*+*τ*_, …, *x*_*t*+(*d*−1)*τ*_) according to their relative order. The probability distribution of the *d*! possible patterns is then estimated as *P* = {*p*(*π*) : *π* ∈ Π}. This encoding is robust to noise, invariant under monotonic transformations, and ideal for capturing nonlinear dynamics in physiological recordings because it quantifies the temporal structure of the signal through the relative ordering of neighboring values rather than their absolute amplitudes [17].

Two representations of every neural recording were subjected to this analysis. In the first, ordinal patterns were calculated directly from the filtered signal (Figure 1D-top), capturing the temporal organization of signal amplitude and reflecting variations in waveform variability and structure that might suggest changes in network excitability or neuronal synchronization. The second highlighted the timing relationships of oscillatory activity and changes in temporal coordination within neural networks by applying ordinal patterns to the instantaneous phase time series obtained via the Hilbert transform (Figure 1D-bottom). Comparing these two representations provides complementary information: phase-derived patterns isolate the signal’s oscillatory structure, whereas amplitude-derived patterns reflect waveform morphology. This distinction is particularly important when oscillatory timing contains the physiologically significant information.

**Figure 1:**
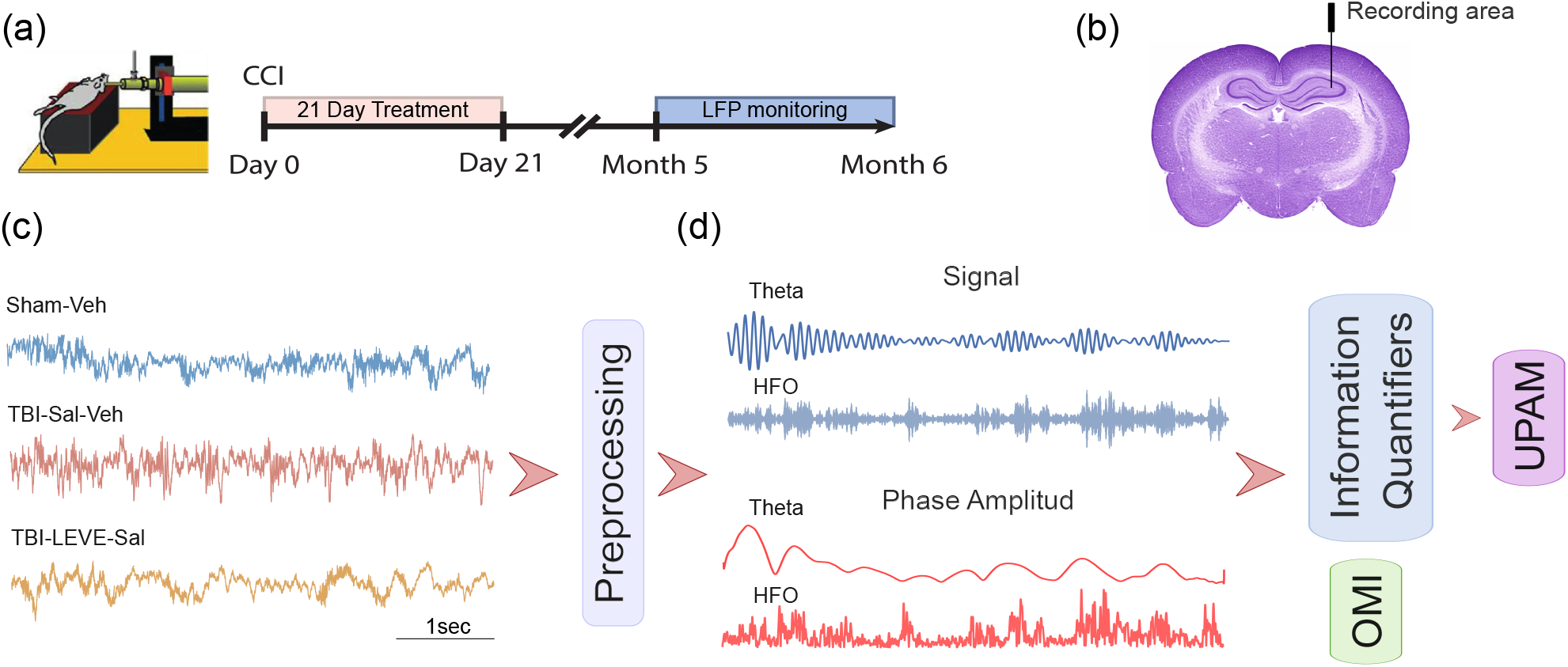
Experimental protocol and signal processing pipeline. **(a)** Timeline of the experimental design: traumatic brain injury (TBI), followed by 21 days of pharmacological treatment with Levetiracetam (LEVE). Five months post-TBI, intracerebral electrodes were implanted, and LFP recordings were performed weekly over four consecutive weeks. **(b)** Anatomical location of the recording electrode in the dorsal hippocampus. **(c)** Representative LFP traces for each experimental group, illustrating the raw signal and its decomposition into the theta (4–8 Hz) and HFO (80–250 Hz) frequency bands, together with the corresponding Hilbert-derived instantaneous amplitude envelopes, used as input for the information-theoretic analysis.

#### 2.4.3 Information Quantifiers

From the ordinal pattern distribution *P*, the following quantifiers were computed to characterize different aspects of the signal dynamics.

##### Permutation Entropy

Permutation Entropy (PE) [13] uses Shannon entropy to measure the diversity of an ordinal pattern distribution to quantify the degree of randomness in a time series:

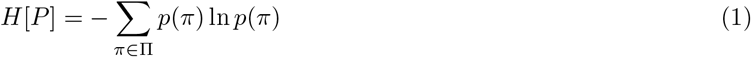

normalized as ℋ = *H*[*P*]*/* ln(*d*!), so that ℋ ∈ [0, 1]. ℋ = 1 corresponds to a uniform distribution, where all *d*! patterns are equally likely (*p*(*π*) = 1*/d*!, ∀ *π* ∈ Π), while ℋ = 0 reflects a fully deterministic system in which a single pattern occurs with probability one (*p*(*π*_*i*_) = 1 and *p*(*π*_*j*_) = 0, ∀ *j≠ i*).

##### Statistical Complexity

Statistical Complexity (SC) [14] quantifies how well a system balances structural organization and randomness. It is defined as:

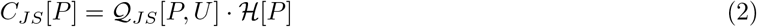

where ℋ [*P*] is the normalized permutation entropy and 𝒬_*JS*_[*P, U*] is the Jensen–Shannon divergence between the observed distribution *P* and the uniform distribution *U*, normalized by its maximum value 𝒬_0_:

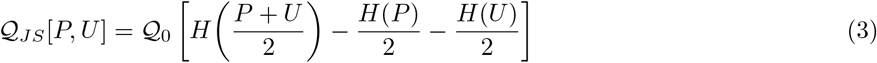

*C*_*JS*_ reaches its maximum for signals with intermediate structure, being zero for both fully random (e.g., white noise) and fully periodic processes, thus allowing discrimination between purely stochastic and organized but complex dynamics.

##### Fisher Information

Fisher Information (FI) [31] quantifies the information an observable random variable carries about an unknown parameter. For ordinal pattern distributions, we employ the discrete formulation proposed in [15]:

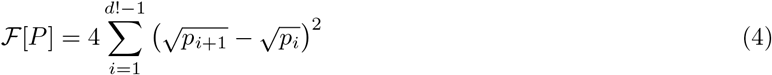

where the sum runs over consecutively ordered patterns *π*_*i*_, is a normalization constant ensuring *F* [*P*] ∈ [0, 1]. Unlike entropy-based measures, FI is sensitive to the “local” structure of the distribution: it increases when probability mass is concentrated in adjacent patterns, reflecting temporal regularity, and decreases for smooth, spread-out distributions associated with irregular dynamics.

##### Permutation Lempel–Ziv Complexity

Permutation Lempel–Ziv Complexity (PLZC) [16] combines the ordinal pattern representation with the Lempel– Ziv compression algorithm [32]. The time series is first transformed into a symbolic sequence *S* = (*π*_1_, *π*_2_, …, *π*_*M*_) by assigning an integer label to each ordinal pattern. The Lempel–Ziv complexity *c*(*S*) is then computed as the number of distinct substrings encountered during a sequential parsing of *S*, and the normalized PLZC is given by:

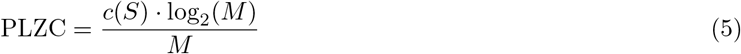

where *M* = *N* − (*d* − 1)*τ* is the length of the symbolic sequence. Unlike the entropy- and divergence-based quantifiers above, PLZC measures algorithmic complexity directly from the symbolic sequence, making it particularly sensitive to the emergence of novel patterns and to abrupt dynamical changes over time.

To better characterize the dynamical properties of the signals, the information quantifiers were analyzed jointly using information planes: the Statistical Complexity–Permutation Entropy (SC–PE) [33], Fisher Information– Permutation Entropy (FI–PE) [34], and Permutation Lempel–Ziv Complexity–Permutation Entropy (PLZC–PE) planes [35]. Representing results in these planes provides a richer description of the underlying dynamics than considering each quantifier in isolation, as different dynamical regimes — purely random, periodic, or structured but complex — occupy distinct regions of each plane.

Together, these three planes form a multidimensional framework to characterize neural dynamics, allowing the identification of differences in randomness, structural organization, local sensitivity, and generative complexity across physiological and pathological conditions.

#### 2.4.4 Dimensionality Reduction and Clustering

Beyond pairwise comparisons in information planes, we aimed to obtain a global view of the dynamical structure across all recordings and conditions. We applied Uniform Manifold Approximation and Projection (UMAP) [18] separately to the information quantifiers derived from the signal amplitude and from the instantaneous phase. In each case, the recordings were represented as feature vectors that include the PE, SC, FI, and PLZC values, yielding a high-dimensional descriptor of the dynamical state of each recording.

UMAP is a nonlinear dimensionality reduction technique that preserves both the local and global topological structure of the data, projecting the high-dimensional feature space onto a two-dimensional embedding that facilitates visual exploration of the data distribution. Unlike linear methods such as principal component analysis (PCA), UMAP captures complex nonlinear relationships between observations, making it particularly suited for neurophysiological data where dynamical differences between conditions may not be linearly separable.

To identify structure within the UMAP embeddings, we applied Hierarchical Density-Based Spatial Clustering of Applications with Noise (HDBSCAN) [19] to the projected data. HDBSCAN is a density-based clustering algorithm that does not require a predefined number of clusters and is robust to noise, assigning observations to clusters based on local density estimates while labelling low-density points as outliers. The algorithm was configured with a minimum cluster size of 3 and a minimum number of samples of 2, allowing the detection of small but cohesive groups within the embedding. The resulting clusters were used to assess whether recordings from different conditions occupy distinct regions of the feature space, providing an unsupervised characterization of the dynamical states captured by the ordinal pattern quantifiers.

#### 2.4.5 Ordinal Modulation Index (OMI)

Cross-frequency interactions were quantified using an Ordinal Modulation Index (OMI), combining phase-amplitude coupling analysis with ordinal pattern encoding [20]. The signal *x*(*t*) was band-pass filtered into a slow component (5–7 Hz, theta band) and a high-frequency component (80–200 Hz, HFO band). The instantaneous phase *ϕ*(*t*) of the slow band was obtained via the Hilbert transform. The fast signal was symbolized using ordinal patterns of embedding dimension *d* = 4 and delay *τ* = 1, yielding a sequence of permutations *π* ∈ Π_*d*_. The phase *ϕ*(*t*) was discretized into *k* = 2 uniform bins, and for each bin *k* the conditional ordinal distribution *P*_*k*_(*π*) was estimated over all time points falling within that phase interval, with *p*_*k*_ denoting the fraction of total time spent in bin *k*. The conditional entropy was then computed as:

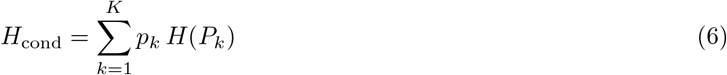

and normalized to define the ordinal modulation index:

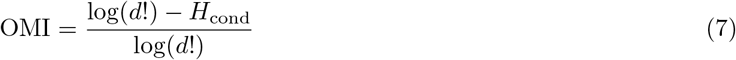

OMI = 0 indicates that the ordinal pattern distribution of the fast signal is uniform across all phase bins – that is, no phase-amplitude coupling is present — while OMI = 1 corresponds to maximal coupling, where the fast signal’s temporal structure is fully determined by the phase of the slow oscillation.

All information-theoretic quantifiers and the Ordinal Modulation Index (OMI) were computed for each 10-second epoch and subsequently averaged across all epochs, yielding a single representative value per recording session.

## 3 Results

### Raw signal analysis

The first analysis focused on information-theoretic quantifiers carried out on raw LFP signals within the theta (4–8 Hz) and HFO (80–200 Hz) frequency bands. The signals were analyzed over the four weeks of the study by taking the average of the information-theoretic metrics from each weekly recording session. We used an embedding dimension of *d* = 4 to discretize the ordinal pattern, and we calculated the best time delay for each band on its own. The optimal delays were *τ*_*θ*_ = 10 for the theta band and *τ*_HFO_ = 1 for the HFO band (see Materials for additional details regarding the techniques employed).

Figure 2 shows the information-plane analyses of LFP signals in the theta frequency band for all experimental groups (Figure 2A). Within the SC–PE plane, both Statistical Complexity (SC) and Permutation Entropy (PE) are reduced in TBI groups relative to sham. The TBI-LEV group displays the lowest SC and PE values across all conditions, a difference that reaches statistical significance compared to sham (Mann–Whitney U test, *p* = 0.038). In the FI–PE plane, the sham group presents significantly lower Fisher Information (FI) and higher PE than all TBI groups (Mann–Whitney U test, *p* = 0.041), with TBI-SAL and TBI-LEV both exhibiting elevated FI values relative to sham. In the PLZC–PE plane, the TBI-LEV group shows the lowest values for both Permutation Lempel–Ziv Complexity (PLZC) and PE, while the sham group exhibits significantly greater values for both measures compared to TBI-SAL and TBI-LEV groups.

**Figure 2:**
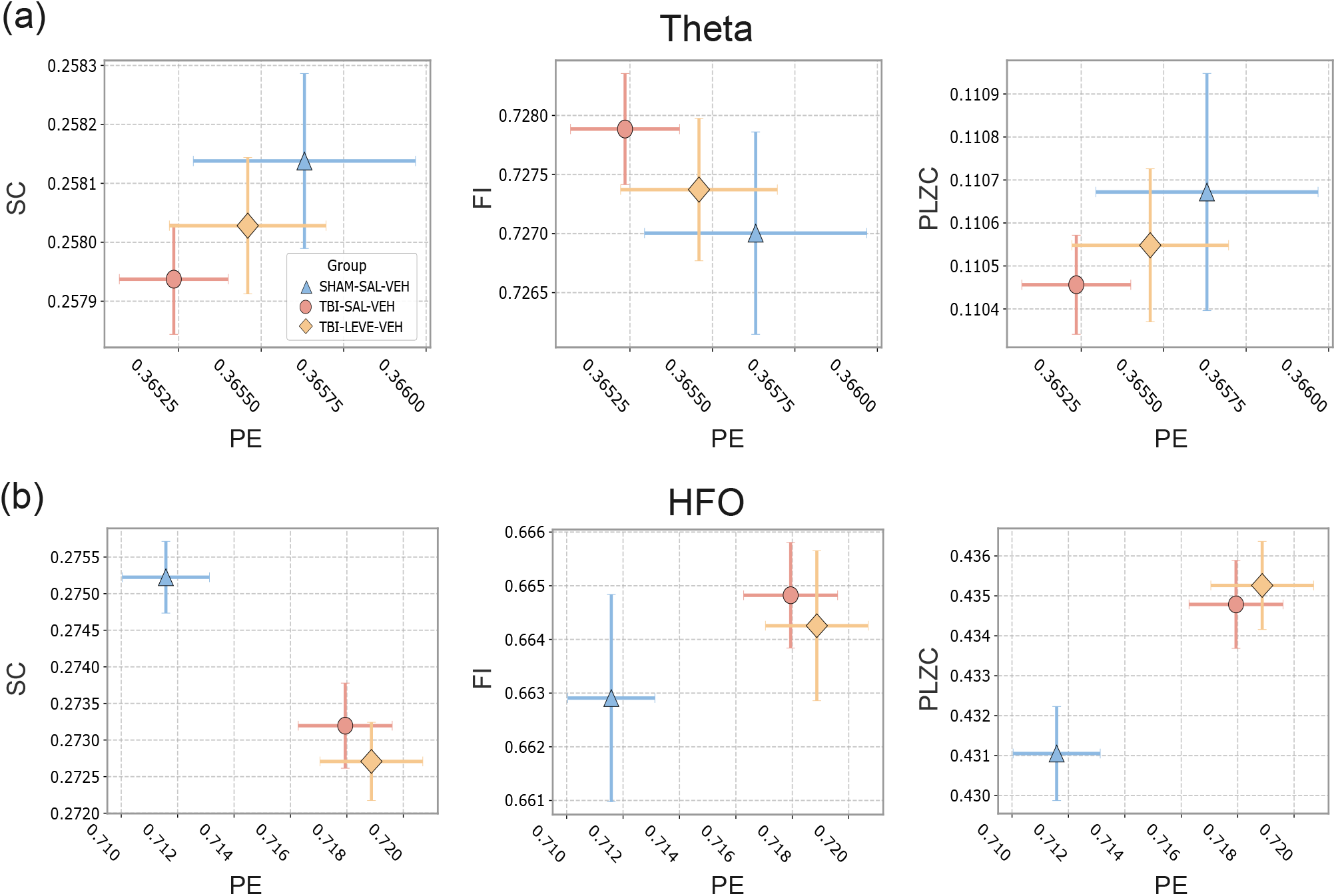
Four-week averaged information-theoretic quantifiers computed from LFP signals across experimental groups (SHAM, TBI-SAL, and TBI-LEV). **(a)** Theta-band (4–8 Hz) analysis represented in the SC–PE, FI–PE, and PLZC–PE information planes. **(b)** HFO-band (80–200 Hz) analysis represented in the same information planes. Each point denotes the group mean, and error bars indicate the standard deviation. SHAM (*n* = 5), TBI-SAL (*n* = 20), TBI-LEV (*n* = 16). Analysis parameters: embedding dimension *d* = 4, time delays *τ*_*θ*_ = 10 and *τ*_HFO_ = 1.

In the HFO band (Figure 2B), within the SC–PE plane, the sham group is clearly separated from all TBI groups, exhibiting higher complexity and lower entropy (Mann–Whitney U test, *p* = 0.033). In TBI animals, the TBI-LEV group shows a small decrease in complexity and a corresponding increase in entropy relative to TBI-SAL, though this difference does not reach statistical significance. In the FI–PE plane, a clear separation between sham and TBI groups is again observed. Within TBI animals, the TBI-LEV group shows a small increase in both FI and PE compared to TBI-SAL. In the PLZC–PE plane, TBI groups exhibit significantly elevated complexity and entropy relative to sham (Mann–Whitney U test, *p* = 0.037). Within TBI animals, the TBI-LEV group shows higher complexity and entropy compared to TBI-SAL.

To analyze the temporal evolution of LFP dynamics, information-theoretic quantifiers were calculated separate for each weekly recording session, employing the same ordinal pattern parameters as utilized in the previous analysis. Figure 3 shows the results for the theta band.

**Figure 3:**
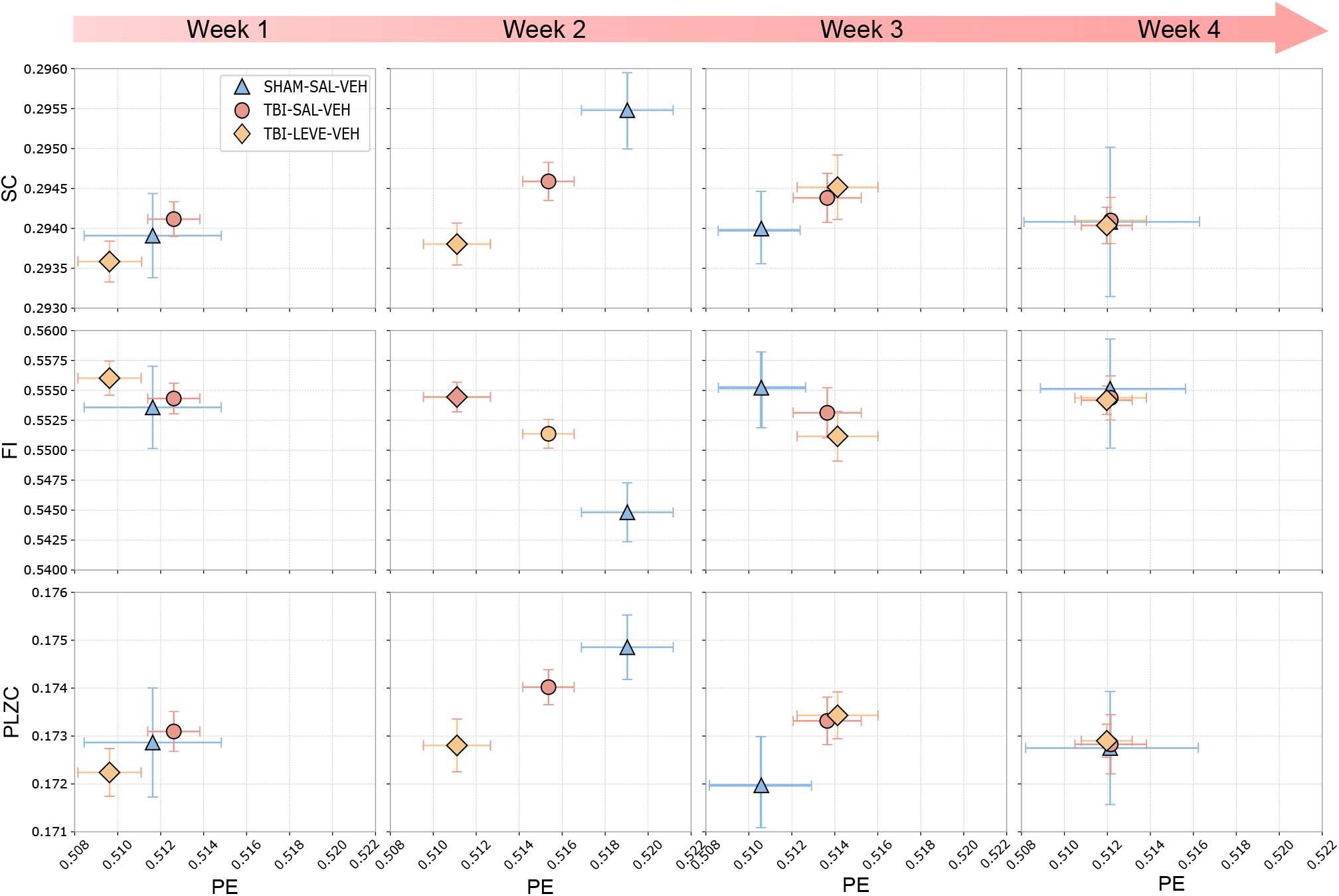
Weekly evolution of information-theoretic quantifiers computed from theta-band (4–8 Hz) LFP signals across experimental groups (SHAM, TBI-SAL, and TBI-LEV). Each row corresponds to a different information plane, from top to bottom: SC–PE, FI–PE, and PLZC–PE. Each point denotes the group mean and error bars indicate the standard deviation. SHAM (*n* = 5), TBI-SAL (*n* = 20), TBI-LEV (*n* = 16). Analysis parameters: embedding dimension *d* = 4, time delay *τ*_*θ*_ = 10

The most significant between-group differences appear at week 2 across all three information planes, whereas weeks 1, 3, and 4 exhibit little to no discernible separation. At week 2 in the SC–PE plane, the sham group presents the highest complexity and entropy values, followed by TBI-SAL, with TBI-LEV showing the lowest values. In the FI–PE plane, TBI-LEV exhibits the highest FI and lowest PE among all groups, while sham displays lower FI and higher PE than both TBI groups. In the PLZC–PE plane at week 2, the sham group surpasses both TBI groups in complexity and entropy, most notably relative to TBI-LEV. At week 3, however, this ordering partially reverses, with sham values falling below those of the TBI groups.

The most significant between-group differences appear at week 2 across all three information planes, whereas weeks 1, 3, and 4 exhibit little to no discernible separation. At week 2 in the SC–PE plane, the sham group presents the highest complexity and entropy values, followed by TBI-SAL, with TBI-LEV showing the lowest values. In the FI–PE plane, a consistent pattern is observed: TBI-LEV exhibits the highest FI and lowest PE among all groups, while sham displays lower FI and higher PE than both TBI groups. In the PLZC–PE plane at week 2, the sham group surpasses TBI groups in both complexity and entropy, most notably relative to TBI-LEV. At week 3, however, this ordering partially reverses, with sham values falling below those of the TBI groups.

Figure 4 displays the results of an analogous week-by-week analysis of HFO-band signals. Unlike the theta band, group differences in the HFO band persist and evolve across weeks 2 through 4, with a consistent pattern of sham–TBI separation gradually emerging.

**Figure 4:**
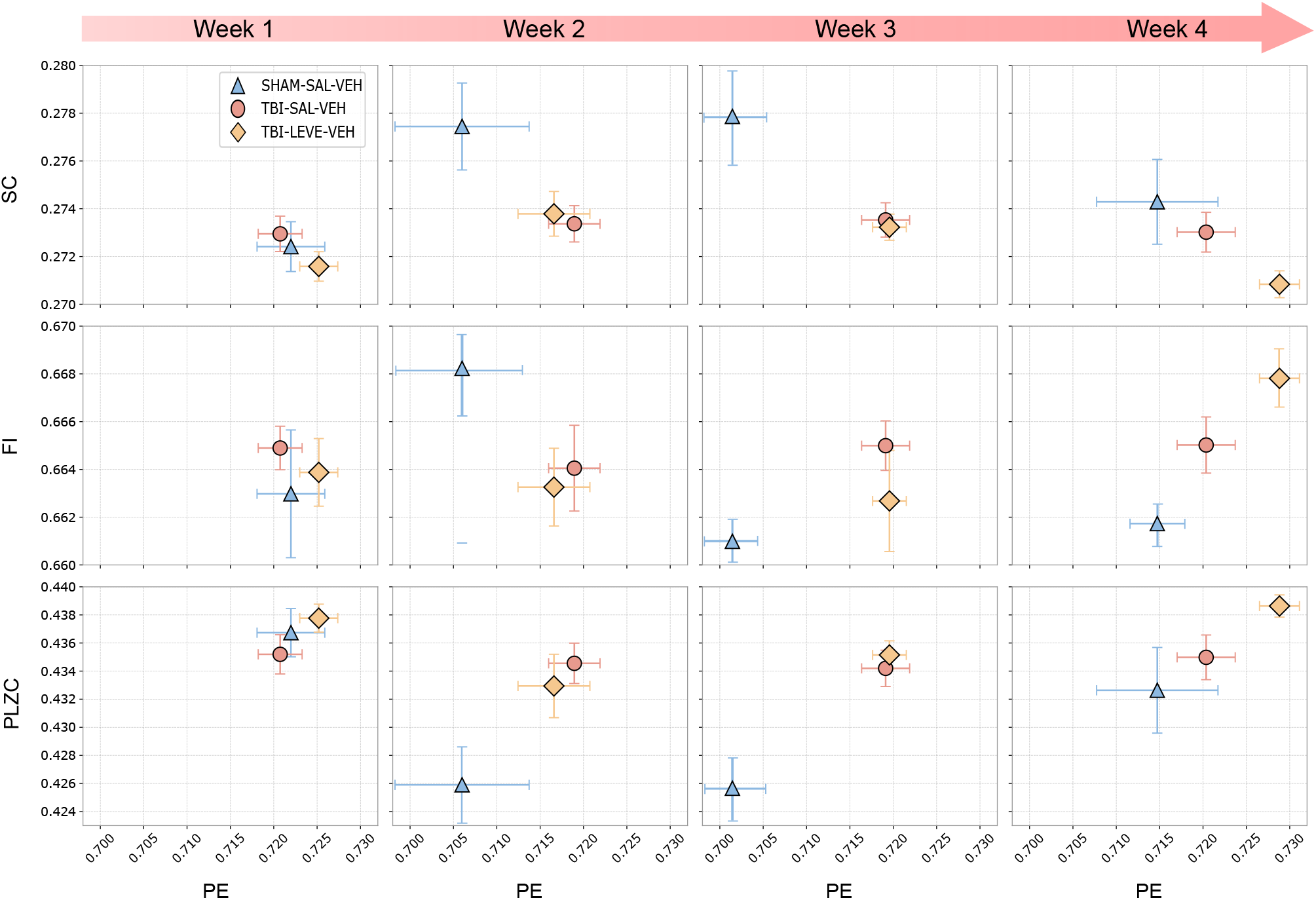
Weekly evolution of information-theoretic quantifiers computed from HFO-band (80–200 Hz) filtered LFP signals across experimental groups (SHAM–SAL–VEH, TBI–SAL–VEH, and TBI–LEVE–VEH). Each row corresponds to a different information plane, from top to bottom: SC–PE, FI–PE, and PLZC–PE. Each point denotes the group mean, and error bars indicate the standard deviation. SHAM (*n* = 5), TBI-SAL (*n* = 20), TBI-LEV (*n* = 16). Analysis parameters: embedding dimension *d* = 4, time delay *τ*_*θ*_ = 10.

In the SC–PE plane, no clear group separation is observed during week 1. Starting at week 2, the sham group shows significantly higher SC and lower PE than both TBI groups. By week 4, the sham and TBI-SAL groups converge, while TBI-LEV continues to exhibit lower complexity and entropy than the other conditions.

In the FI–PE plane, a comparable pattern is observed. Beginning at week 2, the sham group exhibits higher FI and lower PE than both TBI groups. By week 4, two clusters emerge: sham and TBI-SAL form a low-PE/high-FI cluster, while TBI-LEV occupies a high-PE/low-FI cluster.

No between-group differences are observed at week 1 in the PLZC–PE plane. In weeks 2 and 3, the sham group shows a gradual decay of both complexity and entropy, while TBI groups remain largely unchanged from baseline. At week 4, both sham and TBI-LEV show a slight increase of both metrics, indicating a partial recovery. The sham group exhibits a progressive decrease in complexity and entropy during weeks 2 and 3, whereas the TBI groups are relatively stable from baseline. At week 4 both sham and TBI-LEV show a slight increase in both metrics.

### Hilbert Transform Analysis

The second analysis examined information-theoretic quantifiers that were extracted using the Hilbert transform from the instantaneous phase of LFP signals. The same frequency bands as in the previous section were targeted, but the theta band was limited to (5–7 Hz) and the HFO band to (119–161 Hz) due to the mathematical limitations of instantaneous phase estimation, which requires narrowband filtering for reliable phase extraction [36].

Metrics were averaged across the four weekly recording sessions using embedding dimension *d* = 4, with band-specific optimal time delays *τ*_*θ*_ = 20 and *τ*_HFO_ = 1 (see Supplemental Material S3 for methodological details).

The instantaneous phase analysis for both frequency bands in the information-plane is shown in Figure 5. For the theta band (Figure 5A), the sham group displayed higher complexity and entropy compared to TBI-SAL in the SC–PE and PLZC–PE planes (Mann–Whitney U test, *p* = 0.038), but no significant difference compared to TBI-LEV. In the FI–PE plane, the sham group showed lower FI and higher PE than TBI-SAL, with the greatest separation observed relative to the untreated group. In the theta band (Figure 5A), the sham group has higher complexity and entropy than TBI-SAL in the SC–PE and PLZC–PE planes (Mann–Whitney U test, *p* = 0.038), but not with TBI-LEV. The sham group exhibits lower FI and higher PE than TBI-SAL in the FI–PE plane. The greatest separation is observed with respect to the untreated group.

**Figure 5:**
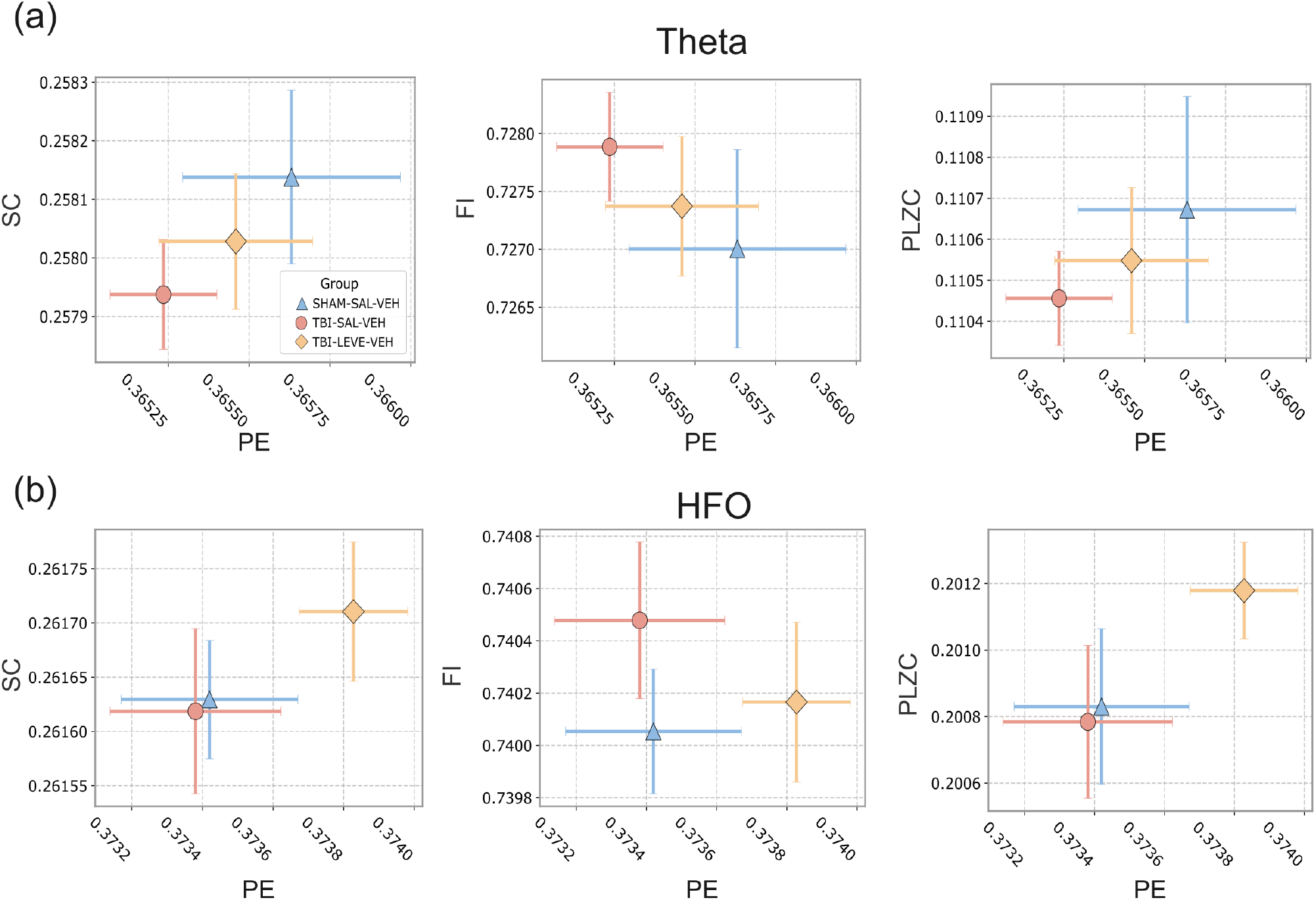
Four-week averaged information-theoretic quantifiers computed from the instantaneous phase of LFP signals across experimental groups (SHAM, TBI-SAL, and TBI-LEV). **(a)** Theta-band (5–7 Hz) analysis represented in the SC–PE, FI–PE, and PLZC–PE information planes. **(b)** HFO-band (119–161 Hz) analysis represented in the same information planes. Each point denotes the group mean, and error bars indicate the standard deviation. SHAM (*n* = 5), TBI-SAL (*n* = 20), TBI-LEV (*n* = 16). Analysis parameters: embedding dimension *d* = 4, time delays *τ*_*θ*_ = 20 and *τ*_HFO_ = 1.

In the HFO band (Figure 5B), the sham and TBI-SAL groups are in a region of lower complexity and entropy in the SC–PE and PLZC–PE planes, while TBI-LEV shifts toward higher values (Mann–Whitney U test, *p* = 0.038). In the FI–PE plane, TBI-SAL has a larger FI and smaller PE than the sham or TBI-LEV, while TBI-LEV has an increasing entropy and decreasing FI.

The week-by-week analysis of the instantaneous phase signals showed limited between-group differentiation, with meaningful separation restricted to week 2 in the theta band (SC–PE and FI–PE planes) and weeks 3–4 in the HFO band (SC–PE and PLZC–PE planes); no consistent temporal structure was observed across the remaining planes or time points.

### UMAP and HDBSCAN Clustering

The information-theoretic quantifiers calculated in the preceding section were divided into four sets for the UMAP analysis: instantaneous phase signals in both bands and raw signals in the theta and HFO bands. Each set was processed with HDBSCAN clustering after UMAP dimensionality reduction.

For both phase amplitude and raw signal analysis, there was no discernible cluster separation between experimental groups. The raw HFO band was an exception, as HDBSCAN found three distinct clusters, one of which was made up only of sham animals (Figure 6A).

**Figure 6:**
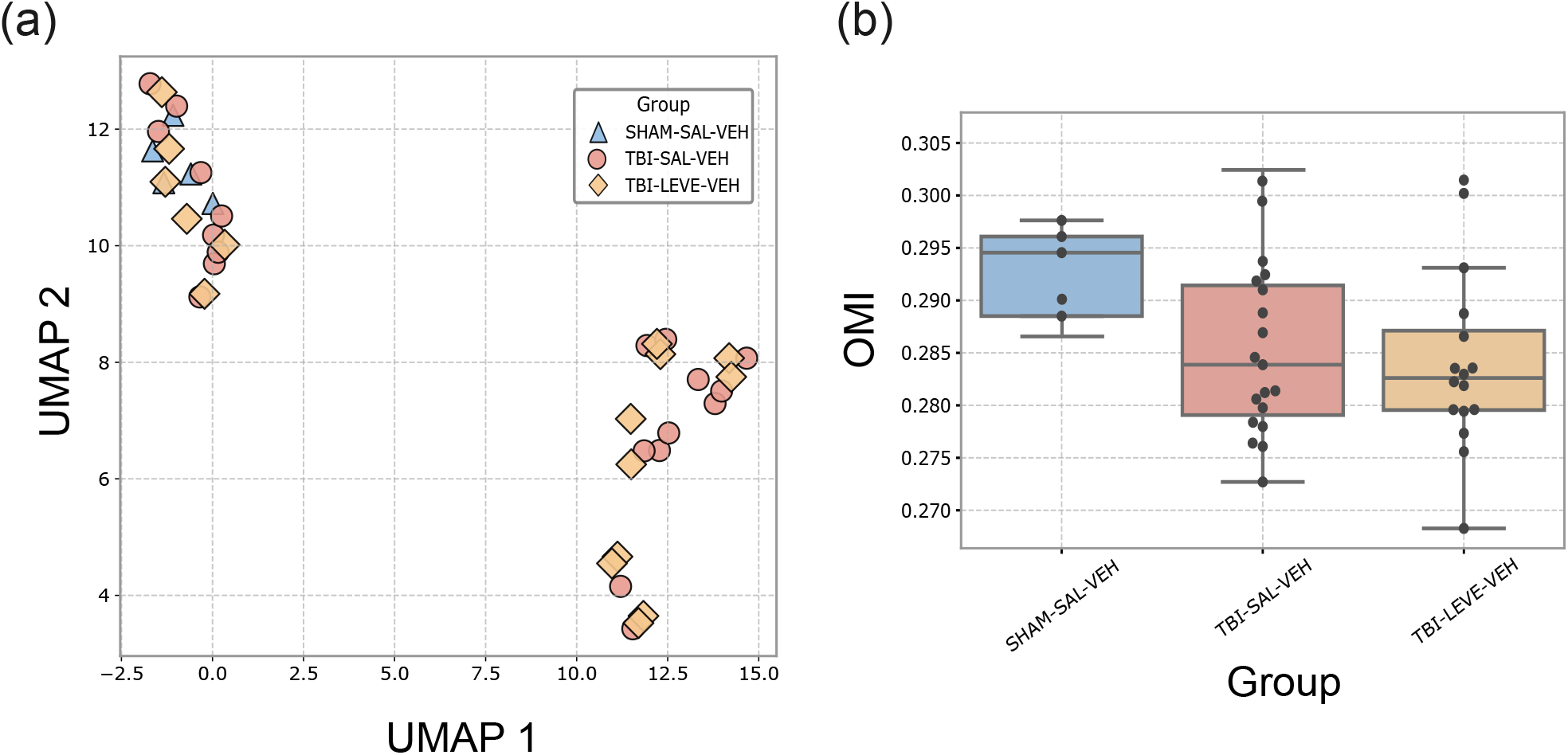
**(a)** UMAP projection based on information-theoretic quantifiers (PE, SC, FI, and PLZC) computed from raw HFO-band (80–200 Hz) LFP signals, with HDBSCAN-identified clusters highlighted. **(b)** Ordinal Modulation Index (OMI) between theta and HFO bands across experimental groups. Analysis parameters: *K* = 20 phase bins for the theta band (5–7 Hz), and embedding dimension *d* = 4 with time delay *τ* = 1 for the HFO band (80–200 Hz).

### Ordinal Modulation Index

For the Ordinal Modulation Index (OMI) analysis, the low-frequency theta band was filtered at 5–7 Hz and the instantaneous phase extracted via the Hilbert transform, discretized into *K* = 20 bins. High-frequency amplitude was obtained from the 80–200 Hz band and discretized using ordinal patterns with parameters *d* = 4 and *τ* = 1.

Figure 6B shows that, regardless of treatment condition, the sham group shows a significantly higher theta– HFO coupling than TBI groups (Mann–Whitney U test, *p* = 0.041). No significant differences were found between the treatment group and the untreated TBI.

## 4 Discussion

In this study, we studied the brain dynamics of mice with traumatic brain injury (TBI) subjected to Levetiracetam drug treatments for 21 days post-injury. The examination of these dynamics was conducted via Local Field Potential (LFP) recordings in the hippocampus of the subjects, carried out weekly over a period of one month, beginning five months post-treatment.

We looked at the LFP signals in two different ways: first, we examined the raw signal filtered in the theta and HFO frequency bands; second, we looked at the amplitude envelope of the phase in those same bands. We used information-entropy planes based on ordinal patterns (SC–PE, FI–PE, and PLZC–PE) to characterize them. We additionally employed UMAP and HDBSCAN, two methods for reducing dimensionality and clustering, on the full set of extracted quantifiers. Lastly, an Ordinal Modulation Index (OMI) algorithm was used to evaluate the relationship between HFO oscillations and the theta band.

### Theta band dynamics

The observed decrease in both complexity and entropy in TBI animals across all information planes (SC–PE, FI– PE, and PLZC–PE) indicates a disruption of hippocampal theta dynamics post-injury. Prior research has reported decreased theta power in hippocampal LFP recordings following TBI, including reduced theta spectral energy in CA1 during exploratory tasks [37], attenuated theta-band firing rhythms at 7 and 14 days post-injury [25], and persistent declines in theta power during spatial navigation [38], coupled with impaired synchronization of CA1 neuronal activity to the prevailing theta oscillation [5]. In the ordinal-pattern framework employed here, such disruption manifests as a reduction of both signal variability and structured complexity—a characterization that transcends spectral power by capturing the nonlinear temporal structure of oscillatory dynamics. Notably, analogous results were obtained from the instantaneous phase signal, indicating that the reduction of dynamic richness in the theta band extends beyond the raw oscillatory signal to its amplitude modulation structure.

The pattern seen in the FI-PE plane gives us more information about what triggered this disruption. A higher Fisher information value means that the signal has more regularity in space and time, which means that it has a more stereotypical dynamic. The sham group, with lower FI and higher PE, shows rich stochastic complexity instead of rigid periodicity. This is a sign of a flexible, high-dimensional oscillatory regime. On the other hand, TBI animals are in a part of the plane with a higher FI and a lower PE, which is consistent with a dynamic that is limited and has little diversity.

In TBI groups, the LEVE-only treatment resulted in the most significant decrease in complexity and entropy, achieving the minimal values across all planes and both signal representations. At first glance, this finding seems to go against reports that levetiracetam increases hippocampal theta power in vitro [39]. But spectral power only shows the linear parts of the signal; ordinal-pattern quantifiers are sensitive to nonlinear temporal dependencies that don’t have to change with power. Consequently, a signal may demonstrate increased spectral energy while concurrently displaying reduced complexity and entropy. This distinction highlights a principal advantage of the information-theoretic approach compared to traditional spectral analysis in characterizing pathological brain dynamics.

### HFO band dynamics

High-frequency oscillations recorded from the hippocampus represent a mixture of physiological and pathological phenomena. Physiological ripples (80–200 Hz) are associated with sharp-wave complexes during memory consolidation and are produced by synchronous inhibitory postsynaptic potentials in interneuron populations. In contrast, pathological high-frequency oscillations (HFOs), including fast ripples exceeding 250 Hz, result from abnormally synchronous bursting of pyramidal cell clusters and are considered biomarkers of epileptogenic tissue. After traumatic brain injury (TBI), an increase in pathological high-frequency oscillations (HFOs) has been consistently observed in hippocampal recordings, with rapid ripple rates indicating which animals will develop post-traumatic epilepsy and indicating the development of a hyperexcitable, hyperconnected network [40, 41]. Our recordings were taken five months after the injury, which is well into the chronic epileptogenic phase. Also, physiological and pathological HFOs can’t be clearly separated from macroelectrode LFP recordings [42, 43]. Because of this, the HFO band signal we looked at here is likely a mix of both types of events. This ambiguity, rather than being a challenge, encourages the utilization of ordinal-pattern quantifiers: instead of using frequency or amplitude thresholds to categorize individual events, the information-entropy planes indicate the overall temporal arrangement of the HFO-band signal, capturing variations in dynamic structure that remain undetectable through spectral analysis alone.

Focus on the analysis of the raw signal. In the SC–PE plane, the sham group is located near the maximum complexity vertex, which is where intermediate permutation entropy meets high statistical complexity. This means that the signal is in a state between order and randomness, which is a state that corresponds to rich, structured dynamics [33]. Both TBI groups are clearly shifted away from this area and toward lower complexity. This means that the HFO-band signal lost its structured nonlinear organization after the injury. The distinction between sham and TBI groups is significantly more evident in the HFO band compared to the theta band, indicating that ordinal-pattern quantifiers are especially responsive to injury-related alterations at elevated frequencies.

The dissociation between SC and PLZC across the complexity–entropy planes is a significant feature of the HFO raw results. While PLZC rises and moves toward higher values in the PLZC–PE plane, SC falls in TBI animals compared to sham animals, indicating a loss of structured complexity. By taking into account the different sensitivities of these two quantifiers within the ordinal-pattern framework, this seeming contradiction is resolved. The degree to which the signal resides in a structured, intermediate dynamical regime is captured by SC, which is derived from the distance between the observed ordinal probability distribution and both the perfectly ordered and perfectly random limits. In contrast, PLZC includes the Lempel–Ziv complexity component, which quantifies the variety of unique ordinal patterns in the signal; a higher PLZC thus corresponds to greater pattern diversity [16]. Therefore, the increase in PLZC in TBI animals is consistent with a signal that gains stochastic rather than organized pattern diversity while losing structured complexity (lower SC)–indicative of a shift toward a more irregular, less coherent dynamical regime. Rather than becoming more periodic, the injured network loses the structured nonlinear organization that characterizes healthy HFO dynamics, replacing it with a higher-dimensional but unstructured variability.

A significantly distinct organizational structure is revealed by the phase amplitude representation, which has unique implications for how treatment effects are interpreted. The pharmacologically treated TBI group are displaced toward significantly higher values in the SC–PE and PLZC–PE planes, whereas the sham and untreated TBI (TBI–SAL–VEH) groups co-locate in an area of lower complexity and entropy. One of the most striking results of the current work is this pattern, which opposes the group structure seen in the raw signal, where Sham was clearly separated from all TBI groups.

The co-location of Sham and TBI-SAL in the SC–PE and PLZC–PE planes reveals that the untreated injured brain and the healthy brain share a similar ordinal-pattern structure at the level of HFO amplitude modulation. This suggests that the complexity and entropy of the amplitude envelope occupy the same region of the information plane, without implying identical dynamics. Leve treatment reorganizes the amplitude modulation dynamics of HFOs, shifting TBI-LEV toward a higher-complexity, higher-entropy regime. In the FI–PE plane, TBI-SAL occupies a region of lower FI and higher PE, consistent with a stochastic, low-structure amplitude modulation, while TBI-LEV exhibits decreased entropy and rearranged local structure. This is geometrically consistent with the SC–PE results: LEVE administration introduces ordinal structure into the amplitude envelope of HFO activity, whereas the untreated TBI phase signal behaves as a high-entropy, low-complexity process.

When combined, the raw and phase amplitude analyses show a dissociation that provides biological and methodological insights. The overall temporal regularity of HFO events–a combination of physiological and pathological oscillations with the injury signature predominating–is captured by the raw signal. The slow modulation of HFO amplitude over time, which is more closely associated with the relationship between high-frequency bursting activity and hippocampal network states, can be detected by the phase amplitude signal. The fact that treatment effects are only discernible in the phase representation highlights the complementary sensitivity of the two signal representations: neither one by itself offers a comprehensive picture of the network reorganization induced by antiepileptic treatment after traumatic brain injury, and their combined analysis—made possible in this case by the ordinal-pattern framework—is necessary to uncover the full structure of the dynamics.

### Longitudinal structure of the recordings

In both frequency bands and signal representations, the first recording session, which took place shortly after the electrodes were put in, always gave quantifier values that were very different from those of the following weeks. Starting in the second week, the values became more stable and were very similar to the four-week averages that were reported in the main analyses. This pattern is most simply explained by the body’s quick response to the insertion of the electrode: in the first few days after implantation, local neuronal circuits undergo mechanical and inflammatory changes that temporarily change LFP dynamics regardless of the experimental condition [44]. The recordings from the first week are considered a stabilization period, and the longitudinal analyses presented here center on weeks two to four.

During this stable window, TBI animals exhibit reduced week-to-week variability in their quantifier values compared to sham animals, whose trajectories exhibit greater fluctuations across sessions. This difference should be interpreted with caution, as it may partially reflect the smaller number of sham animals rather than a true difference in dynamical stability between groups. There is no clear difference from week to week in either the theta or HFO bands when looking at phase amplitude. This suggests that the amplitude modulation structure of the LFP is more stable over the recording period than the raw signal dynamics.

### Theta–HFO coupling: Ordinal Modulation Index

Phase-amplitude coupling (PAC) between low-frequency oscillations and high–frequency activity is a fundamental mechanism of hippocampal network computation. In a healthy hippocampus, the phase of theta oscillations structures the timing of high-frequency events, such as gamma bursts and sharp-wave ripples, thereby offering a temporal framework for information encoding and retrieval [45, 46]. This cross-frequency architecture is significantly compromised subsequent to TBI: Recent laminar recordings in CA1 of injured rats demonstrate significant reductions in theta–gamma phase-amplitude coupling, accompanied by diminished oscillatory power, decreased spike-field coherence, and impaired interneuron entrainment to local oscillations [6]. These results identify PAC as a sensitive indicator of post-TBI circuit dysfunction, signifying the disruption of the inhibitory interneuron networks that regulate hippocampal temporal coding.

In this study, we measured theta-HFO coupling with an Ordinal Modulation Index (OMI), which is a version of the classical Modulation Index of Tort et al. [21] that fits into the ordinal-pattern framework. The OMI quantifies the statistical dependence between theta phase and the ordinal structure of the HFO-band signal, rather than relying on the amplitude of the filtered signal at each theta phase bin. This gives a measure of coupling that is strong enough to handle the nonlinear and non-stationary nature of in vivo LFP recordings. The OMI consequently expands the cross-frequency coupling analysis into the information-theoretic framework established in this study, ensuring methodological coherence throughout the entire analytical process.

The main result of the OMI analysis is a significant decrease in theta–HFO coupling in TBI animals compared to the sham group: TBI animals exhibit no significant coupling, while sham animals show a measurable and significant modulation of HFO ordinal structure by theta phase. This result extends the findings to the theta– HFO frequency pair and the chronic post-injury time point investigated here, and it is consistent with the wider disruption of hippocampal cross-frequency architecture reported after TBI [6].

The absence of significant OMI in TBI groups—including those whose information-plane quantifiers approached sham values—suggests that cross-frequency coordination between theta and HFO dynamics remains persistently disrupted even when treatment partially restores the marginal complexity and entropy of individual bands. This dissociation between cross-frequency coupling and within-band dynamics is a notable finding, as it implies that inter-band coordination and local band dynamics may recover independently and that coupling measures may be more sensitive to the long-term sequelae of TBI than single-band quantifiers alone.

From a methodological standpoint, the OMI demonstrates that ordinal-pattern tools can be naturally extended to cross-frequency analysis, complementing the information-entropy planes applied to individual bands.

### Dimensionality reduction and clustering

There was no apparent distinction between the experimental groups when the entire set of ordinal-pattern quantifiers from the HFO raw signal was projected onto a two-dimensional UMAP embedding, and HDBSCAN clustering was applied. Sham animals and a subset of pharmacologically treated TBI animals were found to form a single cluster, with no identifiable structure separating the other groups. The modest between-group differences seen in the individual information planes for this signal representation are consistent with this result, which shows that the HFO raw feature space does not partition cleanly along experimental condition boundaries when all quantifiers are taken into account jointly in an unsupervised setting.

### Limitations and future directions

The current study has a number of limitations that should be noted. First, comparisons involving the healthy control condition may have less statistical power because there are fewer sham animals than the mean group size of the treated TBI cohorts. This calls for careful interpretation of effect sizes involving the sham group. Second, no information is available from the acute or short–term post–injury period because all recordings were made at a single chronic time point, roughly five months after treatment ended. This makes it difficult to describe the ordinal-pattern signatures’ temporal evolution and raises the question of whether the dynamical differences between groups are stable or progressive over the epileptogenic timeline, as well as when they first appear. Third, it was not possible to distinguish between animals that developed post-traumatic epilepsy and those that did not due to the recording protocol. Individual recording sessions were restricted to about an hour due to the large number of animals and the logistical limitations of the experimental design–a window too small to accurately record spontaneous seizures.

The overall dynamics of a mixed population of potentially epileptic and non-epileptic TBI animals are thus reflected in the ordinal-pattern signatures presented here. Fourth, the dynamical changes found here cannot be directly correlated with functional cognitive outcomes due to the lack of concurrent behavioral readouts; our laboratory is currently working on this.

These limitations will be addressed in multiple ways in future work. To fully characterize the trajectory of hippocampal dynamical reorganization after traumatic brain injury and treatment, longitudinal designs covering the acute, subacute, and chronic post-injury phases will be required. Prospective seizure detection will also be made possible by long-duration recordings with continuous video monitoring. This will enable the ordinal-pattern characterization to be stratified by epileptic outcome and test whether the quantifiers introduced here can function as early biomarkers of post-traumatic epileptogenesis.

On the methodological side, the framework can be extended through the inclusion of additional ordinal-pattern quantifiers — such as weighted permutation entropy and ordinal Rényi entropy — which may offer complementary sensitivity to specific dynamical regimes not fully resolved by the measures employed here. Furthermore, the complete quantifier set could serve as input to supervised classification pipelines, enabling prediction of individual epileptic outcome from chronic LFP recordings and providing a translational bridge between the dynamical characterization developed here and clinical biomarker applications.

## 5 Conclusions

The ordinal-pattern framework applied to hippocampal LFP recordings reveals consistent differences in entropy, complexity, and Fisher information between sham and TBI animals in both the theta and HFO frequency bands across raw and phase amplitude signal representations. These changes reflect a persistent reorganization of hippocampal network dynamics in the chronic post-injury period that is detectable through information-theoretic quantifiers. In addition, theta–HFO coupling as measured by the Ordinal Modulation Index is significantly reduced in all TBI animals with or without medication, relative to sham, indicating a disruption of cross-frequency coordination that is not recovered by the antiepileptic treatments used. Together, these results support the use of ordinal-pattern quantifiers as a sensitive and interpretable tool for characterizing pathological hippocampal dynamics following traumatic brain injury.

## Supplemental Data

### Data Availability Statement

The data supporting the findings of this study are available from the corresponding author upon reasonable request.

