## supplemental for "Analysis of Long-Term Neuronal Dynamics via Ordinal Pattern Quantifiers Following Traumatic Brain Injury and Pharmacological Modulation"

#### S1 Automatic Segment Quality Control

Continuous LFP signals were segmented into non-overlapping windows of 10 s. Let $x(t)$ a segment of length be $N$ sampled at frequency $f_s$. Each segment was evaluated using spectral, temporal, and statistical criteria.

**Frequency-domain SNR.** Power spectral density (PSD) was estimated using Welch's method with $n_{\text{perseg}} = \min(N/2, 2f_s)$. A frequency-domain signal-to-noise ratio ($\text{SNR}_{\text{freq}}$) was computed as:

$$\text{SNR}_{\text{freq}} = \frac{\langle P(f) \rangle_{1-100\,\text{Hz}}}{\langle P(f) \rangle_{>100\,\text{Hz}}}$$

where $\langle P(f) \rangle$ denotes the mean PSD within the specified frequency band. Segments with $\text{SNR}_{\text{freq}} < 15$ were rejected.

**Time-domain SNR.** A robust time-domain signal-to-noise ratio was defined as:

$$\text{SNR}_{\text{time}} = \frac{\text{median}(|x|)}{\text{IQR}(x)}$$

where IQR denotes the interquartile range. Segments with $\text{SNR}_{\text{time}} < 0.5$ were rejected.

**Variance criterion.** Segment variance $\sigma^2$ was computed and compared against the mean variance across all segments from the same channel. Segments with variance exceeding this mean value were discarded.

**Hurst exponent.** The Hurst exponent $H$ was estimated using a log–log regression of the standard deviation of lagged differences:

$$\tau(\ell) = \sqrt{\text{std}(x_{t+\ell} - x_t)}$$

for lags $2 \leq \ell \leq \min(100, N/2)$. $H$ was obtained as the slope of the linear fit between $\log(\ell)$ and $\log(\tau(\ell))$. Segments were retained only if $H$ fell within the adaptive range defined by the 10th–90th percentiles of the Hurst distribution across segments of the same channel.

**Final classification.** A segment was considered valid only if it satisfied all four criteria ($SNR_{freq}$, $SNR_{time}$, variance, and Hurst exponent). Rejected segments were classified according to the first violated criterion.

### S2 Ordinal Modulation Index (OMI)

To quantify cross-frequency interactions between slow oscillations and high-frequency activity, we employed an ordinal-based modulation index (OMI), inspired by the classical phase–amplitude coupling framework [? ] and permutation entropy methods [? ].

##### S2.0.1 Signal preprocessing

The raw signal $x(t)$ was band-pass filtered into:

- a slow component $x_\ell(t)$ in the theta band (5–7 Hz),
- a fast component $x_h(t)$ in the high-frequency oscillation (HFO) range (80–200 Hz).

Filtering was performed using zero-phase Butterworth filters to avoid phase distortions.

The instantaneous phase of the slow component was obtained via the Hilbert transform:

$$\phi(t) = \arg\left(\mathcal{H}[x_\ell(t)]\right), \tag{S1}$$

where $\mathcal{H}[\cdot]$ denotes the analytic signal.

##### S2.0.2 Phase discretization

The phase $\phi(t) \in [-\pi, \pi)$ was discretized into $K$ equally spaced bins:

$$B_k = \left[-\pi + \frac{2\pi(k-1)}{K}, -\pi + \frac{2\pi k}{K}\right), \quad k = 1, \ldots, K. \tag{S2}$$

Each time point $t$ was assigned to a phase bin $k$ such that $\phi(t) \in B_k$.

##### S2.0.3 Ordinal pattern encoding

The fast signal $x_h(t)$ was symbolized using ordinal patterns following the Bandt–Pompe method [? ]. For each time $t$, embedding vectors were constructed as:

$$\mathbf{v}(t) = \left[x_h(t), x_h(t+\tau), \ldots, x_h(t+(D-1)\tau)\right], \tag{S3}$$

where $D$ is the embedding dimension and $\tau$ is the delay.

Each vector $\mathbf{v}(t)$ was mapped to a permutation $\pi \in \Pi_D$, where $\Pi_D$ is the set of all $D!$ possible ordinal patterns.

##### S2.0.4 Conditional distributions

For each phase bin $k$, we estimated the conditional distribution of ordinal patterns:

$$P_k(\pi) = P(\Pi = \pi \mid \phi(t) \in B_k). \tag{S4}$$

The corresponding Shannon entropy was computed as:

$$H_k = -\sum_{\pi \in \Pi_D} P_k(\pi) \log P_k(\pi). \tag{S5}$$

The phase-averaged conditional entropy was defined as:

$$H_{\text{cond}} = \sum_{k=1}^{K} p_k H_k, \tag{S6}$$

where $p_k$ denotes the relative frequency of samples in bin $k$.

##### S2.0.5 Ordinal Modulation Index

The OMI was defined as the normalized deviation from the maximum entropy:

$$\text{OMI} = \frac{H_{\max} - H_{\text{cond}}}{H_{\max}}, \tag{S7}$$

where

$$H_{\max} = \log(D!) \tag{S8}$$

is the entropy of the uniform distribution over ordinal patterns.

##### S2.0.6 Statistical validation

To assess statistical significance, surrogate data were generated by circularly shifting the phase time series $\phi(t)$ relative to the ordinal pattern sequence, thereby preserving the temporal structure of both signals while destroying their alignment.

For each recording, $N = 200$ surrogate realizations were generated, and the corresponding $\text{OMI}_{\text{surr}}$ values were computed.

A $p$-value was estimated as:

$$p = \frac{1}{N} \sum_{i=1}^{N} \mathbb{I}\left(\text{OMI}_{\text{surr}}^{(i)} \geq \text{OMI}_{\text{real}}\right), \tag{S9}$$

where $\mathbb{I}(\cdot)$ is the indicator function.

Additionally, a $z$-score was computed as:

$$z = \frac{\text{OMI}_{\text{real}} - \mu_{\text{surr}}}{\sigma_{\text{surr}}}, \tag{S10}$$

where $\mu_{\text{surr}}$ and $\sigma_{\text{surr}}$ denote the mean and standard deviation of the surrogate distribution.

### S3 Bandwidth selection for Hilbert-based phase analysis

To ensure a meaningful estimation of instantaneous phase using the Hilbert transform, signals must satisfy the narrowband condition, whereby the amplitude envelope varies slowly relative to the carrier oscillation [1, 2]. In practice, this requires limiting the bandwidth of the filtered signal around a central frequency $f_c$.

Based on this criterion, the bandwidth $\Delta f$ was constrained to remain sufficiently small relative to the center frequency:

$$\Delta f \lesssim \alpha\, f_c, \tag{S11}$$

where $\alpha \in [0.3, 0.5]$ controls the trade-off between temporal resolution and spectral specificity. Narrower bands improve phase interpretability by enforcing quasi-monocomponent behavior, while excessively narrow filters may distort temporal dynamics or reduce signal-to-noise ratio.

This heuristic is consistent with established guidelines for time-frequency analysis and phase estimation, which emphasize that reliable instantaneous phase requires signals with limited spectral spread and well-defined dominant frequency components [1, 2, 3].

Accordingly, for Hilbert-based analyses we employed narrower frequency bands compared to those used in raw signal analysis, ensuring that the extracted phase signals satisfied the narrowband assumption required for valid phase-amplitude coupling estimation.

Applying this criterion to the theta band, with central frequency $f_c = 6$ Hz and $\alpha = 0.5$, the resulting bandwidth was $\Delta f_\theta = 3$ Hz, yielding the range (5–7 Hz).

For high-frequency oscillations (HFO), using $f_c = 140$ Hz and $\alpha = 0.3$, the estimated bandwidth was $\Delta f_{\mathrm{HFO}} = 42$ Hz, corresponding to the interval (119–161 Hz).

### S4 Selection of optimal delay parameter ($\tau_{\mathrm{opt}}$)

The embedding delay $\tau$ determines the temporal scale at which ordinal patterns are constructed. For a given embedding dimension $d$, the delay effectively sets the temporal span of the embedding vector, $(d-1)\tau$, and thus must be matched to the intrinsic timescale of the signal. For band-limited signals, this intrinsic timescale is governed by the dominant frequency content, such that slower oscillations require larger delays, whereas faster oscillations require smaller delays.

Based on this relationship, an initial range of candidate delays was defined from the sampling frequency $f_s$ and the frequency band of interest $[f_{\min}, f_{\max}]$:

$$\tau_{\min} \sim \frac{f_s}{k\, f_{\max}}, \quad \tau_{\max} \sim \frac{f_s}{k\, f_{\min}}, \tag{S12}$$

where $k$ represents the number of samples per oscillatory cycle required to adequately resolve the dynamics ($k \in [4, 10]$). This formulation reflects the inverse relationship between frequency and temporal scale, ensuring that the embedding captures meaningful ordinal structure without introducing redundancy (for excessively small $\tau$) or loss of dynamical information (for excessively large $\tau$).

Within this frequency-informed range, the optimal delay $\tau_{\mathrm{opt}}$ was determined using information-theoretic measures derived from the ordinal pattern distribution [4]. Due to the finite bandwidth of the signals, variability in the data, and noise, the optimization typically yielded a plateau or multiple local extrema rather than a single sharp optimum. Therefore, $\tau_{\text{opt}}$ was defined as a range of delays over which the information measures remained stable and near-optimal, consistent with established approaches in nonlinear time-series analysis and delay embedding [5, 6].

Applying this frequency-informed criterion with $k = 6$, we empirically determined the optimal delay ranges for each signal type and frequency band. For raw band-pass filtered signals in the theta range (4–8 Hz), the optimal delays were found within $\tau_\theta \in [10, 42]$, with best performance observed at $\tau_\theta = 10$. In the case of high-frequency oscillations (HFO, 80–200 Hz), the optimal delays were restricted to $\tau_{\text{HFO}} \in [1, 2]$, with $\tau_{\text{HFO}} = 1$ yielding the most consistent results.

For Hilbert-transformed signals, where the theta component was defined in the narrower band (5–7 Hz), the optimal delay range shifted to $\tau_\theta \in [20, 34]$, with $\tau_\theta = 20$ providing the best performance. Similarly, for the HFO amplitude component (119–161 Hz), the optimal delays remained in the range $\tau_{\text{HFO}} \in [1, 2]$, with $\tau_{\text{HFO}} = 1$ showing the highest discriminative power across experimental conditions.

This scale-dependent behavior is consistent with previous studies highlighting the importance of matching embedding parameters to the dominant temporal structure of the signal, particularly in neural time series [3].
